# A cellular assay for spike/ACE2 fusion: quantification of fusion-inhibitory antibodies after COVID-19 and vaccination

**DOI:** 10.1101/2022.06.09.495433

**Authors:** Fabien Abdul, Pascale Ribaux, Aurélie Caillon, Astrid Malézieux-Picard, Virginie Prendki, Nikolay Zhukovsky, Flavien Delhaes, Karl-Heinz Krause, Olivier Preynat-Seauve

## Abstract

Not all antibodies against SARS-CoV-2 inhibit viral entry and hence infection. Neutralizing antibodies are more likely to reflect real immunity, however certain of these tests investigate protein/protein interaction rather than the fusion event. Viral and pseudoviral entry assays detect functionally active antibodies, however they are cumbersome and burdened by biosafety and standardization issues. We have developed a Spike/ACE2-dependant cell-to-cell fusion assay, based on a split luciferase. Hela cells stably transduced with Spike and a large fragment of luciferase were co-cultured with Hela cells transduced with ACE2 and the complementary small fragment of luciferase. Within 24h, cell fusion occured allowing the measurement of luminescence. Light emission was abolished in the absence of Spike and reduced in the presence of an inhibitor of Spike-processing proteases. Serum samples from COVID-19-negative, non-vaccinated individuals, or sera from patients at the moment of first symptoms did not lead to a significant reduction of fusion. In contrast, sera from COVID-19-positive patients as well as sera from vaccinated individuals reduced the fusion. In conclusion, we report a new method measuring fusion-inhibitory antibodies in serum, combining the advantage of a functional full Spike/ACE2 interaction with a high degree of standardization, easily allowing automation in a standard bio-safety environment.

## INTRODUCTION

Only a small subset of the overall antibodies produced against SARS-CoV-2 are neutralizing ^1^ This fraction, namely neutralizing antibodies (nAbs), is crucial because binds to viral antigens in a manner that reduce viral infectivity, in contrast to “non-nAbs” that are not protective. A large number of individuals worldwide have acquired a post-infection or vaccine-induced seroconversion against SARS-CoV-2. A majority of infected patients develop a seroconversion but with individual variations in antibody levels^2, 3^ whereas mRNA vaccines always induce seroconversion^4, 5^. However, age, cancer and immunosuppression are sometimes associated with lower levels of antibodies after infection or vaccination^6, 7^ Importantly, seroconversion does not reflect protective immunity and a report showed that 12% of sera from infected patients containing antibodies do not possess significant levels of nAbs^8^. Other studies reported nAbs in 98% of the infected individuals after six months^9^, with a persistence in 89%–97% of individuals after one year^10–12^ After mRNA vaccination, nAbs have been reported to persist at least six months after the second dose^1^. The pandemic evolution coupled to the introduction of vaccines have imposed a selection pressure causing the evolution of mutants^13^. These mutants possess one or more mutations in their Spike that prevent the neutralization by antibodies generated by the previous strain. It has been reported a reduced neutralizing activity of beta-induced antibodies to delta^12^ Also, sera from vaccinated patients reported a 40-fold reduction of neutralizing activity against omicron^14^ Regarding these multiple parameters influencing the humoral immunity against SARS-CoV-2, it is impossible to predict if an individual is currently protected against a defined variant. Hence, assays that can detect nAbs at both individual and community levels must be developed. An ideal test for mass testing should be rapid, cheap, reproducible, automatable and performed with minimal safety precautions. Numerous analytical methods have been developed to measure anti-SARS-CoV-2 nAbs in the serum.

As the Receptor-Binding Domain (RBD) of the viral Spike was identified to be crucial for entry, it was initially suggested that antibodies against RBD could reflect nAbs. Several immunoassays were developed to detect anti-RBD antibodies^15, 16^. The surrogate virus neutralization test is a competitive ELISA where the nAbs compete with an in vitro immobilized and quantifiable RBD / ACE2 interaction. Various kits are available and are 17-20 21 sensitive and specific^17–20^, but fail to detect nAbs at low levels and suffer from a high false positive rate due to the wrong detection of non-nAbs^22^ Unfortunately for these RBD-based assays, anti-RBD are not necessarily neutralizing^23^ and nAbs targeting epitopes outside the RBD are known, such as the N-terminal domain of S1^24^ and the S2 domain^25^. The plaque reduction neutralization test is the gold standard: a fixed load of a living virus is exposed to the serum prior the infection of cultured cells and counting of cytopathic effects (the formed plaques). It is a manual, labour intensive and time-consuming procedure (72–96 h) requiring a biosafety level 3 and some strains do not produce plaques. The counting of infected cells and the time of analysis can be improved by including a reporter gene into the viral genome^26, 27^ but it still requires level 3 environment. Pseudotyped viruses that are replication-incompetent, express a reporter gene and use the same entry mechanism as SARS-CoV-2 have been developed^28^. They require at least 48h of infection and can be performed under more acceptable bio-safety conditions (level 2).

In vitro cell fusion between cells expressing a viral protein and cells expressing its receptor is a method used for the quantification of virus entry in target cells. The rationale of this approach is the formation of syncytia in some virus-infected tissues, induced by the fusion of infected cells with neighbouring cells. Thus, syncytia are multinucleated enlarged cells induced by the surface expression of viral proteins that interact with their receptors expressed in neighbouring cells, creating a fusogenic event and allowing virus spreading without the need of endocytosis. As syncytia were described in the lungs of COVID-19 patients^29^, in vitro cell fusion assays between spike-expressing cells and ACE2-expressing cells has been described and used for the study of chemical inhibitors of the Spike/ACE2 interaction^30–32^ With the goal to target all the epitopes of the full-length Spike with a low cost, a high rapidity of execution and a high level of standardization, we have developed a cell fusion assay emitting luminescence in a Spike/ACE2 interaction-dependant manner. As luminescence is reduced by the presence of antibodies inhibiting the Spike/ACE2 interaction, it allows a sensitive and specific quantification of inhibitory antibodies in serum. This new assay is rapidly executed (24h), reproducible, cheap, automatable and easily standardized for large scale analyses.

## MATERIAL AND METHODS

### Cells and reagents

Hela cells were cultured in DMEM medium with 4.5 g/l of glucose (Gibco), supplemented with 1% of penicillin and streptomycin, non-essential amino-acids and 1mM sodium pyruvate and 10% of foetal bovine serum (Thermofisher). The used luciferase substrate for the detection of luminescence was The NanoGlo live assay, from Promega.

### Blood samples

Individuals were recruited among the STRAT-CoV and GEROCOVID cohorts, including patients hospitalized for COVID-19 at the Geneva University Hospitals, Switzerland, SARS-CoV-2 infection was diagnosed with a positive Reverse Transcriptase – Polymerase Chain Reaction against SARS-CoV-2 in naso-pharyngeal swabs. Blood samples were collected at admission) and about two weeks later. This study also included volunteers, sampled at different time points depending of the disease and vaccinations with mRNA vaccines. Patients with psychiatric disorders or for whom consent could not be obtained were excluded. Ethics approval was granted by the Cantonal Ethics Research Committee of Geneva, Switzerland (GEROCOVID: no. 2020-01248, Dr V. Prendki and Dr A. Malézieux-Picard, April 2020-May 2021; STRAT-CoV: no. 2020-01070, Dr S. Baggio, Dr N. Vernaz, February 2020 – February 2022).

### Molecular biology

The ACE2-expressing Hela cells (Hela-ACE2) were established from a cDNA ORF purchased from GenSript. The pCG1_SCoV2-S plasmid encoding the original Wuhan Spike (S_wuhan_) (provided by Prof. Dr. Stefan Pöhlmann, University Göttingen, Göttingen, Germany) was used to generate the S-expressing Hela cells (Hela-S). ACE2 and S cDNA ORFs were cloned into pCDH-CMV-MCS-EF1α-Puro using a standard cloning method. Four different variants of S were also generated with the indicated mutations: S_D614G_ (D614G), S_delta_ (E156-F157del, R158G, L452R, T478K, D614G, P681R, D950N), S_lambda_ (G75V, T67I, R246-G252del, D253N, L452Q, F490S, D614G, T859N) and S_omicron_ (A67V, H69-V70del, T95I, G142-V143-Y144del, Y145D, N211del, L212I, G339D, S371L, S373P, S375F, K417N, N440K, G446S, S477N, T478K, E484A, Q493R, G496S, Q498R, N501Y, Y505H, T547K, D614G, H655Y, N679K, P681H, N764K, D796Y, N856K, Q954H, N969K, L981F).

The plasmids SmBiT-PRKACA and LgBiT-PRKAR2A containing respectively the small BiT and the large BiT part of the luciferase were purchased from Promega (NanoBiT® PPI MCS Starter System CAT.# N2014). SmBiT-PRKACA and LgBiT-PRKAR2A cDNA ORFs were cloned into pCDH-CMV-MCS-EF1α-RFP and pCDH-CMV-MCS-EF1α-CopGFP respectively using standard cloning methods.

For recombinant-lentivirus production, lentivector plasmids were transfected in HEK 293T cells using the calcium phosphate method. Hela-S and Hela-ACE2 were established from lentiviral cotransduction resulting in two cell lines expressing respectively Spike/GFP/LgBiT and ACE2/RFP/SmBiT.

### Cell fusion assay

A mixture of 10 000 clonal Hela-S and 10 000 clonal Hela-ACE2 were cocultured in 96-well flat clear bottom white polystyrene microplates (Corning) in 100μl of culture medium in the presence of patient’s serum or control sera at various dilutions (1/8, 1/32, 1/128 and 1/512). After 24h of culture at 37°C under 5% of CO_2_, the medium was removed and cells were rinsed once with HBSS buffer with Calcium and magnesium (Thermofisher) prior addition of 100 μl of the same HBSS buffer. Twenty-five microliters of the substrate diluted at 1/20 in buffer (NanoGlo live assay, Promega, see the manufacturer’s instructions) was added extemporaneously. The microplate was shaked for 1min at 500 rpm, then centrifugated 1min at 300 g prior to the addition of an adhesive white opaque film at the bottom of the plate and luminescence was measured with a spectra-L-luminometer (Molecular devices). Experiments were performed in triplicates. To prevent variability between operators, assays and reagents, the same internal control made of pooled sera from SARS-CoV-2 – negative patients were systematically included. The inhibition was normalized for each serum dilution with this internal control and calculated as the ratio between the luminescence from patient’s serum and the luminescence from the control serum.

## RESULTS

### Coculture of Hela cells expressing the SARS-CoV-2 Spike and Hela expressing ACE2 induces cell fusion

Hela cells were stably transduced with the original Wuhan Spike sequence (S_wuhan_) or some selected variants: The European variant with the D614G mutation (S_D614G_), lambda variant (S_lambda_), delta variant (S_delta_) and omicron variant (S_omicron_). Another Hela cell was stably transduced with the ACE2 receptor, also under the control of a ubiquitous promoter. To allow quantification of fusion, a dual split reporter system has been also introduced in Hela-Spike and Hela-ACE2 lines. The Hela-Spike were co-transduced with a construct expressing a large part of the luciferase, namely the large BiT luciferase (LgBiT) and the Green Fluorescent protein (GFP) whereas Hela-ACE2 were co-transduced with a construct coexpressing the complementary resting small part of luciferase, namely small BiT luciferase (SmBiT)^1^ and the Red Fluorescent Protein (RFP). Upon fusion, the expressed LgBiT and SmBiT recombine and produce a functional luciferase emitting quantifiable luminescence in the presence of a cell-permeable substrate (figure 1A). Cultured alone in a 96 well plate, clonal Hela-S or clonal Hela-ACE2 developed towards a monolayer of single cells (figure 1B,C). In contrast, the coculture induced rapidly (24h later) cells clusters, suggesting fusogenic events and formation of syncytium-like structures (figure 1D). These clusters were double-fluorescent for GFP and RFP, whereas Hela-S alone were GFP+ / RFP- and Hela-ACE2 were GFP-/RFP+ (figure 1E), confirming cell fusions between the two lines. Expectedly, the replacement of Hela-S_wuhan_ by Hela-S_D614G_, Hela-S_delta_, Hela-S_lambda_ increased fusion because all of these variants have more affinity for ACE2. Indeed, the in vitro fusogenic events were strong enough to create rapidly bigger cell clusters having the tendency to detach from the bottom of the plate (figure 1F). In contrast, Hela-S_omicron_ induced clusters similarly to Hela-S_wuhan_.

**Figure 1.**
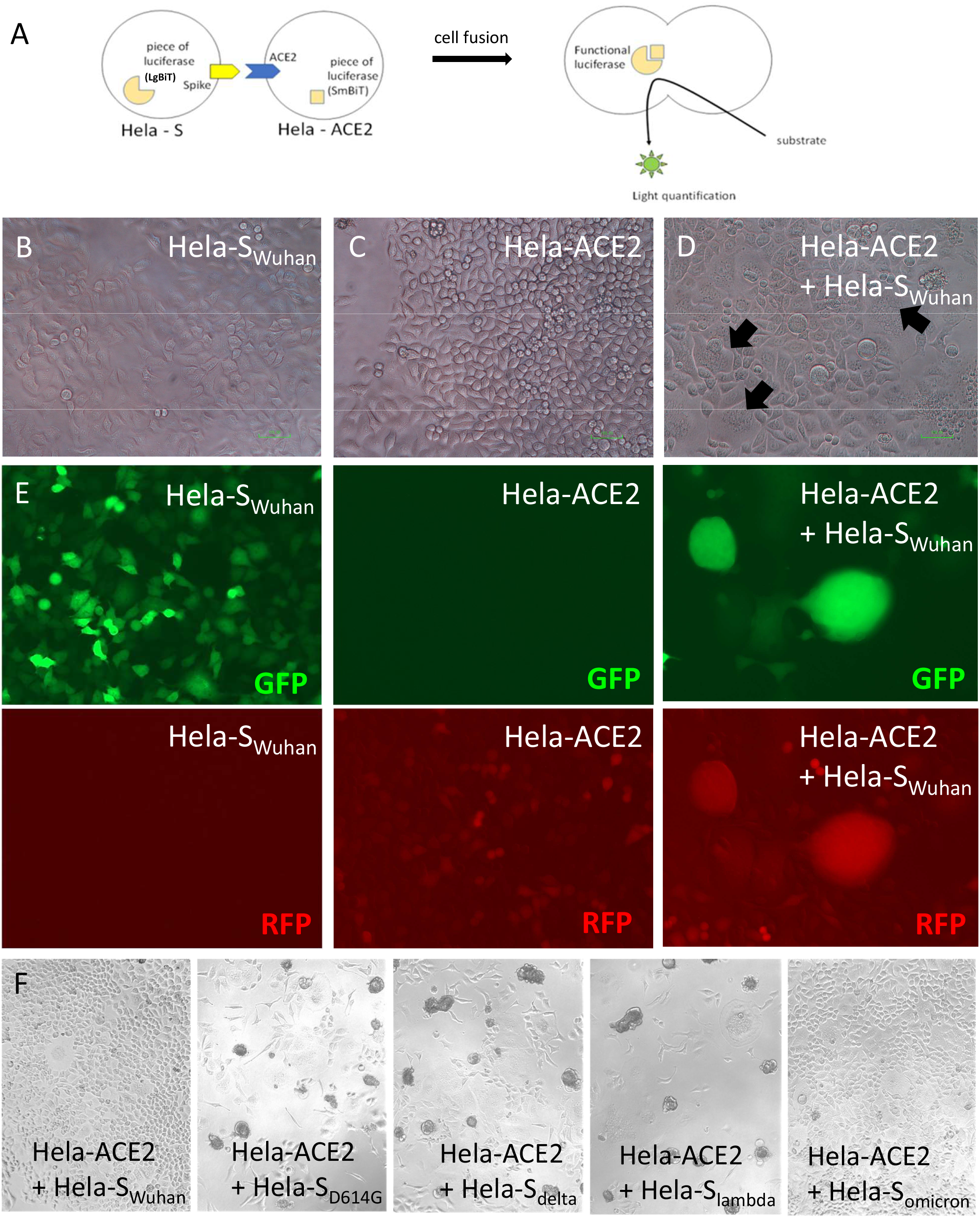
Coculture of Hela-S and Hela ACE2 induces cell fusion. A – Principle of the dual split fusion assay between a Hela-S and a Hela-ACE2. B-F Cells were cultured in 96 well plate for 24h. B, C, D – Microscopic observation of Hela-S (B) or Hela-ACE2 (C) alone versus co-culture (D). The black arrows show cell syncitia indicative of fusion. Magnification x40 E – Observation of the fluorescence emitted by Hela-S or Hela-ACE2 alone versus co-culture. Hela-S were GFP+ and Hela-ACE2 RFP+. Magnification x40 F – Hela cells expressing various variants of Spike were cocultured with Hela-ACE2, prior to a microscopic observation of cells. Magnification x20

With the goal to develop an assay measuring antibodies inhibiting the Spike/ACE2 – dependant cell fusion, serum must be added to the coculture prior to washing and addition of a cell-permeable substrate for luciferase. Washing is mandatory because human serum is known to inhibit the reaction between the luciferase and its substrate. Because detachment of cell clusters, observed with some variants of Spike, must be avoided at the washing step (to ensure reproducibility and use in a large-scale setting), we decided to develop and evaluate an assay using the Hela-S_wuhan_.

Cells cultured alone in a 96 well plate (Hela-S-LgBiT alone or Hela-ACE2-SmBiT alone) did not generate luminescence, in contrast to the co-culture (figure 2A). The signal increased with the cell concentration, confirming a dose-dependent response (not shown). Co-culture using a Hela cell expressing LgBiT but not Spike (Hela-LgBiT) abolished the signal, confirming that the emission of the fluorescence was dependant of the Spike/ACE2 interaction (figure 2A).

**Figure 2.**
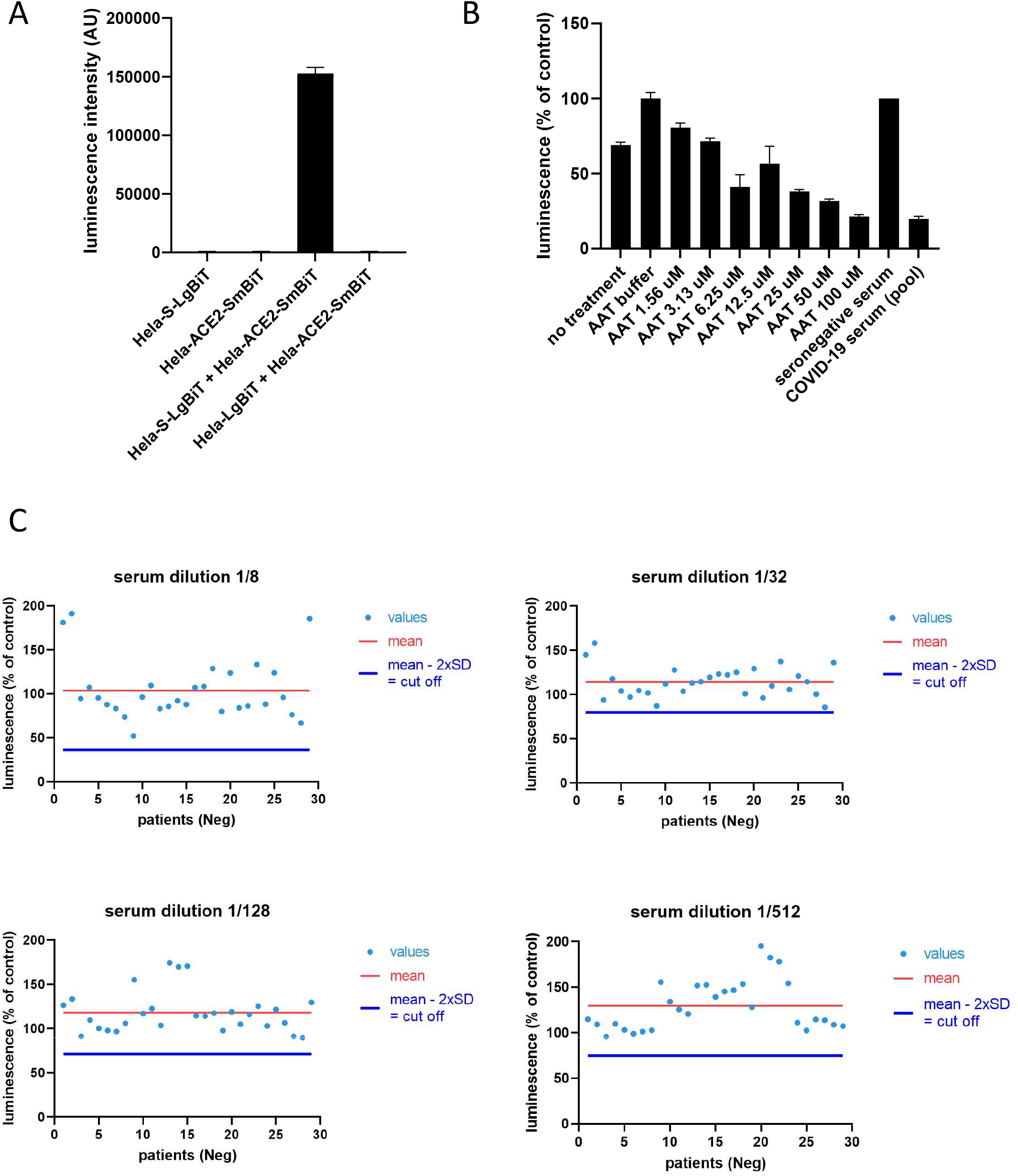
Impact of human seronegative serum on the luminescence emission. A – Various combinations of Hela transduced or not with S, ACE2, LgBiT or SmBiT were cocultured 24h in a 96 well plate, prior to addition of the substrate for luciferase and reading on a luminometer. B – Influence of AAT on cell fusion. Each luminescent signal is normalized with the value in the presence of an internal control, made of another pool of seronegative sera (expressed in %). C – Coculture of Hela-S and Hela-ACE2 in the presence of various dilutions of sera (1/8 to 1/512) from seronegative patients without any COVID-19 history. Each luminescent signal is normalized with the value in the presence of an internal control, made of another pool of seronegative sera (expressed in %).

### Coculture of Hela-S-LgBiT and Hela-ACE2-SmBiT in the presence of various dilutions of seronegative human serum induces emission of a luminescent signal

Because the goal of the assay is to evaluate in human serum the presence of antibodies inhibiting Spike/ACE2-dependant fusion, the impact of human serum on the emission of luminescence was first evaluated. Indeed, factors present in the human serum could interfere with the cellular fusion, independently of immunoglobulins. Several serum dilutions ranging between 1/8 and 1/512 were tested by using samples from 30 seronegative patients without any history of COVID-19 or vaccination (GEROCOVID cohort). To prevent variability between operators, assays and reagents, an internal control made of a pooled seronegative serum was systematically included. Each luminescent signal was then normalized by the internal control and calculated as a ratio (in %). Luminescence emission was proportionally reduced by the addition of increased concentrations of Alpha1 Anti-Trypsin (AAT), an inhibitor of TMPRSS2 protease^33^ (figure 2B) indicating that the fusion was dependant of the processing of Spike and not solely its physical interaction with ACE2. The results from 29 seronegative individuals and for each serum dilution are (figure 2C, associated with the calculation of the average and 2 standard deviations. One patient was considered to be an outlier (probably an unknown seropositivity). The dilution 1/8 showed the higher variability (standard deviation) between patients, suggesting that an excess of serum could impact the reproducibility of the test (figure 2C). The calculation of 2 standard deviations was used to define a cut off value for the detection of a statistically significant reduction of fusion. The value of the cut off was impacted by the serum dilution at 1/8 due to the increased variability, in contrast to higher dilutions. Table 1 shows the numerical values of the average of luminescence (in % of control serum), standard deviation for each serum dilution.

**Table 1.** Average and standard deviation between 29 seronegative sera tested for their impact on the fusion.

| serum<br>dilution | average | standard deviation | 2 standard deviations |
| --- | --- | --- | --- |
| 1/8 | 103.7 | 33.8 | 67.6 |
| 1/32 | 114.1 | 17.1 | 34.1 |
| 1/128 | 118.1 | 23.4 | 46.8 |
| 1/512 | 129.5 | 27.3 | 54.6 |

### Impact of seropositive sera from COVID-19 patients on the emission of a luminescent signal

Fifty-six sera from COVID-19 patients (D614G European variant) were collected at the Geneva University Hospitals, Switzerland, two weeks after the first symptoms, thanks to the STRAT-CoV cohort (Dr V. Prendki and Dr A. Malézieux). A proportion of these sera reduced the normalized emission of luminescence below the cut off of 2 standard deviations defined previously (figure 3A). An effect of the serum dilution was observed: the proportion of samples below the cut off increased from 1/512 to 1/32, with a maximum at 1/32. Thus, compared to seronegative sera, sera from COVID-19 patients were able to reduce the Spike/ACE2 – dependant fusion, suggesting the presence of inhibitory antibodies. We also collected sera from 23 patients (GEROCOVID cohort, Dr V. Prendki and Dr A. Malézieux) at 2 different time points: at the moment of first symptoms of a primo-infection (T1) and 2 weeks later (T2), considering the absence of humoral immunity at T1. These 23 patients were not vaccinated. For a majority of these patients, there was a clear reduction of fusion at T2 compared to T1 (figure 4), confirming that the inhibition was acquired 2 weeks after the first symptoms and not constitutive of the nature of the serum, therefore again suggesting fusion-inhibitory antibodies.

**Figure 3.**
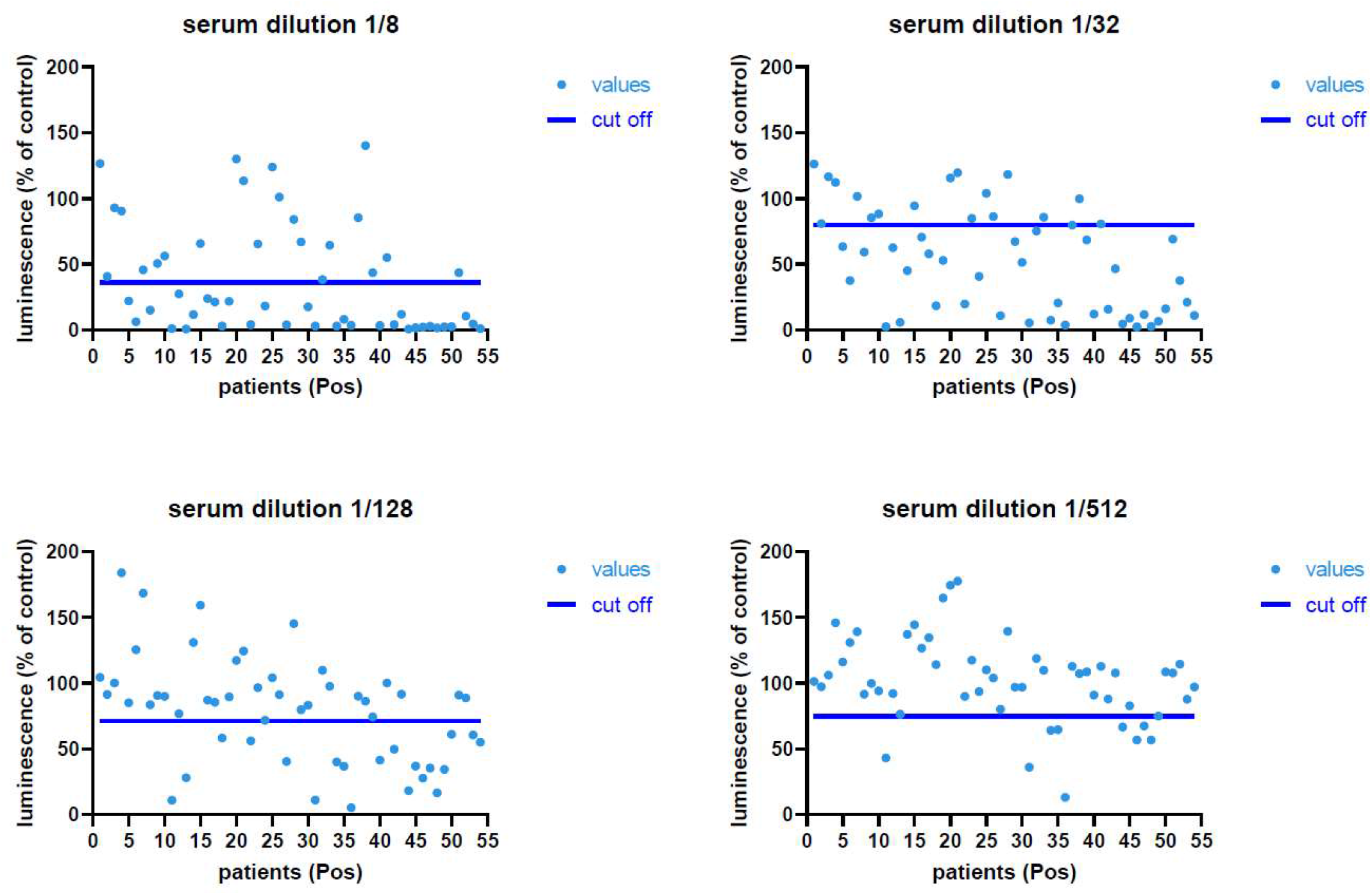
Impact of serum from COVID-19 patients on the luminescence emission. Coculture of Hela-S and Hela-ACE2 in 96 well plate for 24h in the presence of various dilutions of sera (1/8 to 1/512) from COVID-19 patients two weeks after the first symptoms. Each luminescent signal is normalized with the value in the presence of an internal control, made of a pool of seronegative sera (expressed in %).

**Figure 4.**
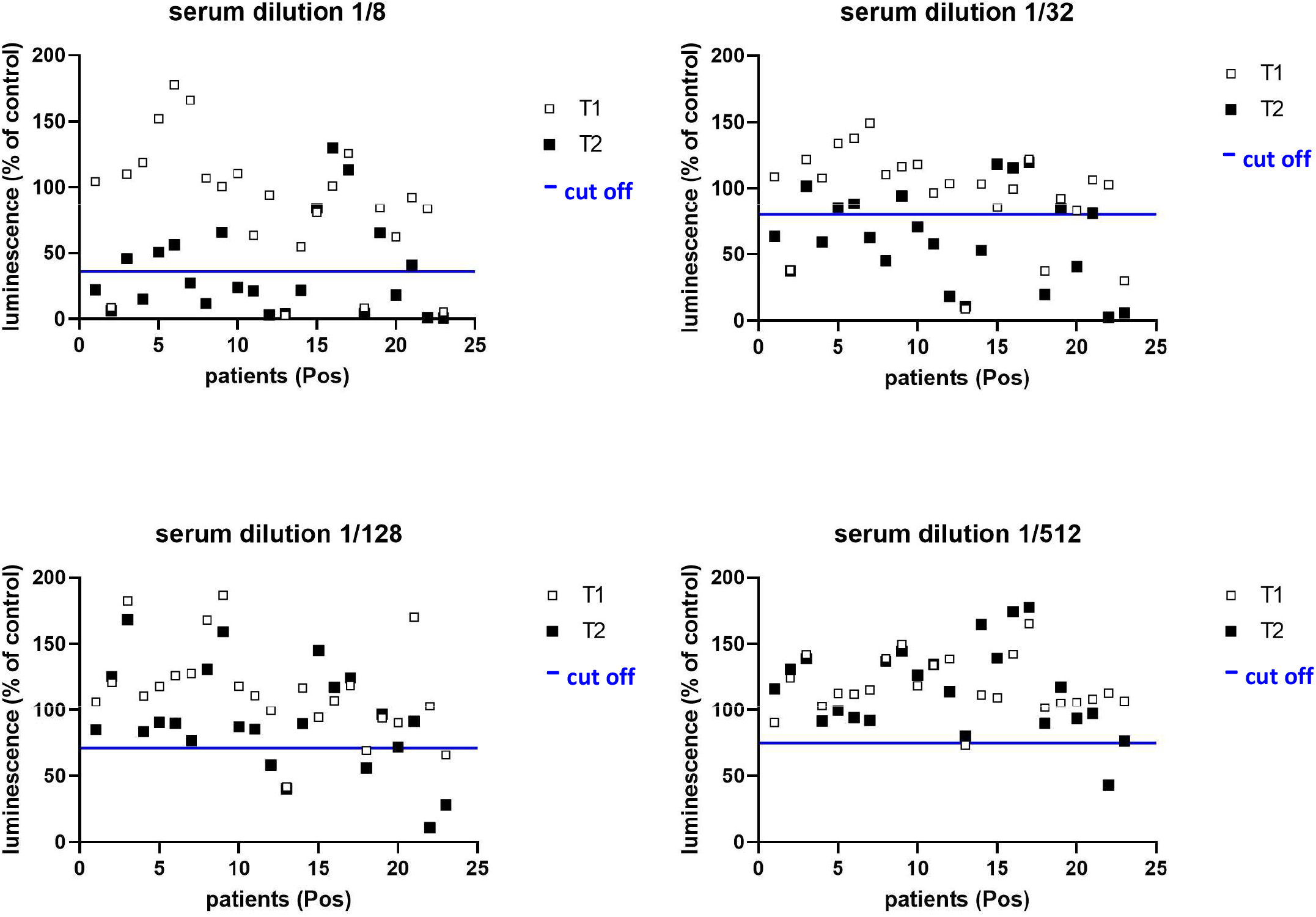
Impact of sera from COVID-19 patients on luminescence emission at the moment of first symptoms and 2 weeks later. Coculture of Hela-S and Hela-ACE2 in the presence of various dilutions of sera (1/8 to 1/512) from COVID-19 patients at the moment of first symptoms (T1) or two weeks after (T2). Each luminescent signal is normalized with the value in the presence of an internal control, made of a pool of seronegative sera (expressed in %).

### Case-reports of the kinetic of fusion inhibitions in vaccinated individuals

Because of the large-scale introduction of vaccination worldwide, it was important to evaluate the inhibition of fusion after vaccination. Figure 5 shows several case-reports. A first individual (ID#1) was primo-infected by the beta variant and did not show any neutralization below the cutoff three weeks after the first symptoms. A mRNA vaccine following primo-infection induced a stable inhibition (below the cutoff) for at least 5 months, maintained by a second mRNA vaccine later. It confirms that fusion-inhibitory antibodies can be detected and followed after mRNA vaccination. Some individuals (ID#2 to ID#5) vaccinated by various series of mRNA vaccines without any previous primo-infection showed stable fusion-inhibitory antibodies after the first vaccination for at least 6 months. On the other hand, other individuals also vaccinated by series of mRNA vaccines (ID#6 to ID#8) only show inhibition after the second dose of vaccine. It is noteworthy that one individual (ID#9) lost fusion-inhibitory antibodies 6 months after the second dose. Supplementary table 1 shows the details of the vaccinal scheme of each individual

**Figure 5.**
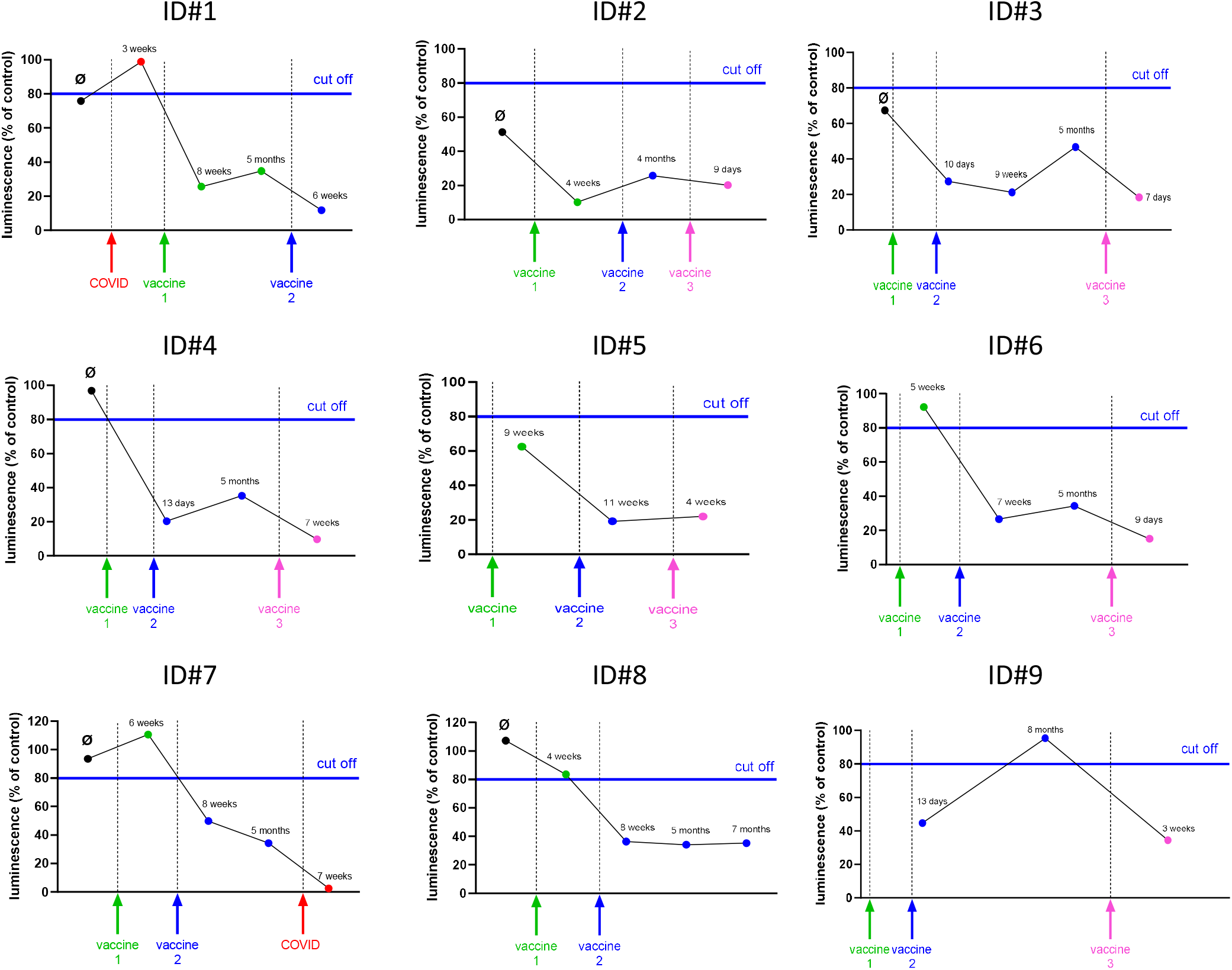
Case-reports of fusion inhibition after mRNA vaccination. Sera (dilution 1/32) from 9 volunteers (ID#1 to ID#9) with various vaccinal schemes were analyzed for their impact on the luminescence emission. Ø = serum before any vaccination/infection. The time of vaccination or infection is indicated by vertical arrows with a color code (red for COVID infection, green for 1^st^ vaccination, blue for 2^nd^ vaccination and pink for 3^rd^ vaccination). The dots correspond to blood sampling and the time laps refer to the delay between blood sampling and the previous vaccination/infection. Each luminescent signal is normalized with the value in the presence of an internal control, made of a pool of seronegative sera (expressed in %).

Together, these observations indicate that the assay can be used for the determination of fusion-inhibitory antibodies following vaccination. As expected, there is a clear variability in the vaccinal response between patients, then reinforcing the need of new assays evaluating the inhibition of the Spike/ACE2 interaction.

## DISCUSSION

Cell-to-cell fusion occurs in vivo because syncytia, which are large multinucleated pneumocytes, are seen in the lungs of COVID-19 patients^34^. Mechanistically, cell-to-cell fusion is similar to virus-cell fusion and thus represents an excellent modelization of viral entry, reinforcing the interest of its use for the detection of inhibitory antibodies. The viral Spike is considered to be the fusogenic molecule. Viral entry begins when the Spike interacts with ACE2. Then Spike is cleaved by TMPRSS2 at the S1/S2 junction site to allow fusion in a zone of the plasma membrane to create an endocytic vesicle engulfing the virus. Viral RNA is transferred to the cytoplasm and translated, producing a Spike protein translocated into the endoplasmic reticulum and transported throughout the Golgi and the membrane. The membrane Spike associates with a ACE2 receptors on neighbouring cells receptor via its RBD domain, and is also again processed by proteases at the S1/S2 sites. S1 domain is released which allows the fusion process between the 2 cells through the formation of a common pore that will expand^34^. Thus, developing an assay measuring inhibitors of fusion will add a more functional evaluation of the impact on viral entry mechanisms rather than only a Spike/ACE2 or RBD/ACE2 physical interaction. Accordingly to this point, we observed that the cell fusion assay performed in the presence of inhibitors of TMPRSS2 reduced fusion, confirming the implication of the molecular machinery of entry. If the Pseudotyped virus entry assay, have also the advantage to modelize the Spike-dependant virus/entry mechanism, the fusion assay has the strong advantage of its reduced cost, simplicity and do not require a bio-safety level 2 environment.

The degree of syncytia formation correlates with the affinity of Spike with ACE2^35, 36^and we observed that changing the original Wuhan Spike in the fusion assay by other variants strongly influenced fusogenicity, with a strong increase seen with the European (D614G), delta and lambda variants. The European variants contains the mutation (D614G) close to the S1/S2 cleavage site, conferring to Spike a more competent binding for ACE2^37, 38^. Pseudovirus assays performed by our group (not shown) and others^39, 40^ have shown that the D614G mutation increases the efficiency of viral entry. Moreover, the D614G mutant produces more syncytia in vitro than the Wuhan strain^36, 41^. In accordance with our observations, the delta variants is more fusogenic than the D614G variant^36, 42, 43^. The impact of the lambda and omicron mutations on fusion are not described precisely and our data shows an impact of lambda, but not omicron.

Because a too strong intensity of fusion induces big clusters detaching from the plate, specific technical adaptations needs to be achieved to assess each variant in the assay. We used the Wuhan sequence in this study as a proof of concept, expectedly exploitable for patients infected by the delta variant and individuals vaccinated with the most current mRNA vaccines. The possibility to adapt easily any variants will represent a strong advantage. The interest in using various Spike mutants in the assay is useful not only for the evaluation of the protective immunity of some populations exposed to defined variants or new mRNA vaccines, but also to understand the immunological cross reactivities between the variants and vaccines.

Another advantage of this fusion assay consists in its rapidity of execution (24h) in multiwell plates, compared to the plaque reduction neutralization test and pseudotyped viruses, moreover in a standard bio-safety environment. It then provides a strong level of reproducibility and standardization adapted for automation and large-scale studies.

There was a clear impact of human serum by itself on fusion, independently of the presence of inhibitory antibodies. Indeed, the increase of serum from seronegative individuals without any history of COVID-19 in the co-culture enhanced the luminescent signal. To explain this phenomenon, we hypothesized a possible proliferation effect of human serum on Hela cells, or the presence of serum proteins facilitating cell-to-cell fusion by an active process or just neutralization of the negative repulsive charges at the surface of cells. However, the use of an internal control made from pooled other seronegative sera, in addition to normalize the different assays, correct this background signal influencing the results.

How to interpret the data to produce numerical values evaluating the neutralization? First, there is the question of the cut off value, that will depend on variants and must be determined in each assay. Two standard deviations below the average of at least 30 seronegative sera appear to be a statistically correct cut off to define the presence of an inhibitor. We observed a clear effect of serum dilution on the obtained value, which was also dependant of each individual. Thus, we would not preconize to select an arbitrary dilution of serum and just render a result in percentage of luminescence compared to the internal control serum. It would represent a risk to overestimate or underestimate the inhibition, because outside of the optimal window. Thus, we more preconize the systematic use of several dilutions and the calculation of the dilution that induce a reduction of 50% of the control signal (namely the half inhibitory concentration or IC50).

## CONCLUSION

We describe in this study a new dual split luciferase-based fusion assay to measure the presence of fusion-inhibitory antibodies in the serum of individuals. It will allow large-scale screening of immunity in populations post disease or vaccination or treatment. This assay is cheap, well standardized for automation and does not require a specific bio-safety environment. Because using a full-length Spike protein and including molecular events mimicking viral entry, this assay can be performed with any variant of Spike or future mRNA vaccines, then providing a new and useful tool to detect antibodies active on fusion.

## ACKNOWLEDGEMENTS

Authors thanks Dr Stéphanie Baggio (Geneva University Hospitals) and Dr Nathalie Vernaz (Geneva University Hospitals) for the availability of the STRAT-CoV cohort of blood samples.

## SUPPLEMENTARY MATERIAL

Table S1. Vaccination scheme of healthy individuals having received mRNA

## AUTHOR CONTRIBUTIONS

Conceptualization, F.A., K.H.K, O.P.S; Methodology, F.A., P.R., A.C., K.H.K, O.P.S; Formal Analysis, F.A., P.R., A.C., K.H.K, O.P.S; Investigation F.A., P.R., A.C., K.H.K, O. P.S; Resources, F.A., P.R., A.C., K.H.K, O.P.S; A.M., V.P., N.Z., F.D.; Data Curation, P. R., A.C., A.M., V.P., O.P.S; Writing – Original Draft Preparation, P.R., O.P.S.; Writing – Review & Editing, F.A., P.R., O.P.S.; Supervision, K.H.K, O.P.S; Funding Acquisition, K.H.K.

## INSTITUTIONAL REVIEW BOARD STATEMENT

The study was approved by the Cantonal Ethics Research Committee of Geneva, Switzerland (GEROCOVID: no. 2020-01248, Dr V. Prendki and Dr A. Malézieux; STRAT-CoV: no. 2020-01070, Dr S. Baggio, Dr N. Vernaz).

## REFERENCES

1. Jeremiah, S. S.; Miyakawa, K.; Ryo, A., Detecting SARS-CoV-2 neutralizing immunity: highlighting the potential of split nanoluciferase technology. J Mol Cell Biol 2022.

2. Dan, J. M.; Mateus, J.; Kato, Y.; Hastie, K. M.; Yu, E. D.; Faliti, C. E.; Grifoni, A.; Ramirez, S. I.; Haupt, S.; Frazier, A.; Nakao, C.; Rayaprolu, V.; Rawlings, S. A.; Peters, B.; Krammer, F.; Simon, V.; Saphire, E. O.; Smith, D. M.; Weiskopf, D.; Sette, A.; Crotty, S., Immunological memory to SARS-CoV-2 assessed for up to 8 months after infection. Science 2021, 371 (6529).

3. Wei, J.; Matthews, P. C.; Stoesser, N.; Maddox, T.; Lorenzi, L.; Studley, R.; Bell, J. I.; Newton, J. N.; Farrar, J.; Diamond, I.; Rourke, E.; Howarth, A.; Marsden, B. D.; Hoosdally, S.; Jones, E. Y.; Stuart, D. I.; Crook, D. W.; Peto, T. E. A.; Pouwels, K. B.; Walker, A. S.; Eyre, D. W.; team, C.-I. S., Anti-spike antibody response to natural SARS-CoV-2 infection in the general population. Nat Commun 2021, 12 (1), 6250.

4. Jackson, L. A.; Anderson, E. J.; Rouphael, N. G.; Roberts, P. C.; Makhene, M.; Coler, R. N.; McCullough, M. P.; Chappell, J. D.; Denison, M. R.; Stevens, L. J.; Pruijssers, A. J.; McDermott, A.; Flach, B.; Doria-Rose, N. A.; Corbett, K. S.; Morabito, K. M.; O’Dell, S.; Schmidt, S. D.; Swanson, P. A., 2nd; Padilla, M.; Mascola, J. R.; Neuzil, K. M.; Bennett, H.; Sun, W.; Peters, E.; Makowski, M.; Albert, J.; Cross, K.; Buchanan, W.; Pikaart-Tautges, R.; Ledgerwood, J. E.; Graham, B. S.; Beigel, J. H.; m, R. N. A. S. G., An mRNA Vaccine against SARS-CoV-2 – Preliminary Report. N Engl J Med 2020, 383 (20), 1920–1931.

5. Walsh, E. E.; Frenck, R. W., Jr.; Falsey, A. R.; Kitchin, N.; Absalon, J.; Gurtman, A.; Lockhart, S.; Neuzil, K.; Mulligan, M. J.; Bailey, R.; Swanson, K. A.; Li, P.; Koury, K.; Kalina, W.; Cooper, D.; Fontes-Garfias, C.; Shi, P. Y.; Tureci, O.; Tompkins, K. R.; Lyke, K. E.; Raabe, V.; Dormitzer, P. R.; Jansen, K. U.; Sahin, U.; Gruber, W. C., Safety and Immunogenicity of Two RNA-Based Covid-19 Vaccine Candidates. N Engl J Med 2020, 383 (25), 2439–2450.

6. Gavriatopoulou, M.; Terpos, E.; Ntanasis-Stathopoulos, I.; Briasoulis, A.; Gumeni, S.; Malandrakis, P.; Fotiou, D.; Migkou, M.; Theodorakakou, F.; Eleutherakis-Papaiakovou, E.; Kanellias, N.; Kastritis, E.; Trougakos, I. P.; Dimopoulos, M. A., Poor neutralizing antibody responses in 106 patients with WM after vaccination against SARS-CoV-2: a prospective study. Blood Adv 2021, 5 (21), 4398–4405.

7. Terpos, E.; Gavriatopoulou, M.; Fotiou, D.; Giatra, C.; Asimakopoulos, I.; Dimou, M.; Sklirou, A. D.; Ntanasis-Stathopoulos, I.; Darmani, I.; Briasoulis, A.; Kastritis, E.; Angelopoulou, M.; Baltadakis, I.; Panayiotidis, P.; Trougakos, I. P.; Vassilakopoulos, T. P.; Pagoni, M.; Dimopoulos, M. A., Poor Neutralizing Antibody Responses in 132 Patients with CLL, NHL and HL after Vaccination against SARS-CoV-2: A Prospective Study. Cancers (Basel) 2021, 13 (17).

8. Chia, W. N.; Zhu, F.; Ong, S. W. X.; Young, B. E.; Fong, S. W.; Le Bert, N.; Tan, C. W.; Tiu, C.; Zhang, J.; Tan, S. Y.; Pada, S.; Chan, Y. H.; Tham, C. Y. L.; Kunasegaran, K.; Chen, M. I.; Low, J. G. H.; Leo, Y. S.; Renia, L.; Bertoletti, A.; Ng, L. F. P.; Lye, D. C.; Wang, L. F., Dynamics of SARS-CoV-2 neutralising antibody responses and duration of immunity: a longitudinal study. Lancet Microbe 2021, 2 (6), e240–e249.

9. Goto, A.; Go, H.; Miyakawa, K.; Yamaoka, Y.; Ohtake, N.; Kubo, S.; Jeremiah, S. S.; Mihara, T.; Senuki, K.; Miyazaki, T.; Ikeda, S.; Ogura, T.; Kato, H.; Matsuba, I.; Sanno, N.; Miyakawa, M.; Ozaki, H.; Kikuoka, M.; Ohashi, Y.; Ryo, A.; Yamanaka, T., Sustained Neutralizing Antibodies 6 Months Following Infection in 376 Japanese COVID-19 Survivors. Front Microbiol 2021, 12, 661187.

10. Epaulard, O.; Buisson, M.; Nemoz, B.; Marechal, M. L.; Terzi, N.; Payen, J. F.; Froidure, M.; Blanc, M.; Mounayar, A. L.; Quenard, F.; Pierre, I.; Pavese, P.; Germi, R.; Grossi, L.; Larrat, S.; Poignard, P.; Lupo, J., Persistence at one year of neutralizing antibodies after SARS-CoV-2 infection: Influence of initial severity and steroid use. J Infect 2022, 84 (3), 418–467.

11. Haveri, A.; Ekstrom, N.; Solastie, A.; Virta, C.; Osterlund, P.; Isosaari, E.; Nohynek, H.; Palmu, A. A.; Melin, M., Persistence of neutralizing antibodies a year after SARS-CoV-2 infection in humans. Eur J Immunol 2021, 51 (12), 3202–3213.

12. Miyakawa, K.; Kubo, S.; Stanleyraj Jeremiah, S.; Go, H.; Yamaoka, Y.; Ohtake, N.; Kato, H.; Ikeda, S.; Mihara, T.; Matsuba, I.; Sanno, N.; Miyakawa, M.; Shinkai, M.; Miyazaki, T.; Ogura, T.; Ito, S.; Kaneko, T.; Yamamoto, K.; Goto, A.; Ryo, A., Persistence of Robust Humoral Immune Response in Coronavirus Disease 2019 Convalescent Individuals Over 12 Months After Infection. Open Forum Infect Dis 2022, 9 (2), ofab626.

13. Krause, P. R.; Fleming, T. R.; Longini, I. M.; Peto, R.; Briand, S.; Heymann, D. L.; Beral, V.; Snape, M. D.; Rees, H.; Ropero, A. M.; Balicer, R. D.; Cramer, J. P.; Munoz-Fontela, C.; Gruber, M.; Gaspar, R.; Singh, J. A.; Subbarao, K.; Van Kerkhove, M. D.; Swaminathan, S.; Ryan, M. J.; Henao-Restrepo, A. M., SARS-CoV-2 Variants and Vaccines. N Engl J Med 2021, 385 (2), 179–186.

14. Cele, S.; Jackson, L.; Khoury, D. S.; Khan, K.; Moyo-Gwete, T.; Tegally, H.; San, J. E.; Cromer, D.; Scheepers, C.; Amoako, D. G.; Karim, F.; Bernstein, M.; Lustig, G.; Archary, D.; Smith, M.; Ganga, Y.; Jule, Z.; Reedoy, K.; Hwa, S. H.; Giandhari, J.; Blackburn, J. M.; Gosnell, B. I.; Abdool Karim, S. S.; Hanekom, W.; Ngs, S. A.; Team, C.-K.; von Gottberg, A.; Bhiman, J. N.; Lessells, R. J.; Moosa, M. S.; Davenport, M. P.; de Oliveira, T.; Moore, P. L.; Sigal, A., Omicron extensively but incompletely escapes Pfizer BNT162b2 neutralization. Nature 2022, 602 (7898), 654–656.

15. Bray, R. A.; Lee, J. H.; Brescia, P.; Kumar, D.; Nong, T.; Shih, R.; Woodle, E. S.; Maltzman, J. S.; Gebel, H. M., Development and Validation of a Multiplex, Bead-based Assay to Detect Antibodies Directed Against SARS-CoV-2 Proteins. Transplantation 2021, 105 (1), 79–89.

16. Kubo, S.; Ohtake, N.; Miyakawa, K.; Jeremiah, S. S.; Yamaoka, Y.; Murohashi, K.; Hagiwara, E.; Mihara, T.; Goto, A.; Yamazaki, E.; Ogura, T.; Kaneko, T.; Yamanaka, T.; Ryo, A., Development of an Automated Chemiluminescence Assay System for Quantitative Measurement of Multiple Anti-SARS-CoV-2 Antibodies. Front Microbiol 2020, 11, 628281.

17. Perera, R.; Ko, R.; Tsang, O. T. Y.; Hui, D. S. C.; Kwan, M. Y. M.; Brackman, C. J.; To, E. M. W.; Yen, H. L.; Leung, K.; Cheng, S. M. S.; Chan, K. H.; Chan, K. C. K.; Li, K. C.; Saif, L.; Barrs, V. R.; Wu, J. T.; Sit, T. H. C.; Poon, L. L. M.; Peiris, M., Evaluation of a SARS-CoV-2 Surrogate Virus Neutralization Test for Detection of Antibody in Human, Canine, Cat, and Hamster Sera. J Clin Microbiol 2021, 59 (2).

18. Putcharoen, O.; Wacharapluesadee, S.; Chia, W. N.; Paitoonpong, L.; Tan, C. W.; Suwanpimolkul, G.; Jantarabenjakul, W.; Ruchisrisarod, C.; Wanthong, P.; Sophonphan, J.; Chariyavilaskul, P.; Wang, L. F.; Hemachudha, T., Early detection of neutralizing antibodies against SARS-CoV-2 in COVID-19 patients in Thailand. PLoS One 2021, 16 (2), e0246864.

19. Tan, C. W.; Chia, W. N.; Qin, X.; Liu, P.; Chen, M. I.; Tiu, C.; Hu, Z.; Chen, V. C.; Young, B. E.; Sia, W. R.; Tan, Y. J.; Foo, R.; Yi, Y.; Lye, D. C.; Anderson, D. E.; Wang, L. F., A SARS-CoV-2 surrogate virus neutralization test based on antibody-mediated blockage of ACE2-spike protein-protein interaction. Nat Biotechnol 2020, 38 (9), 1073–1078.

20. Taylor, S. C.; Hurst, B.; Charlton, C. L.; Bailey, A.; Kanji, J. N.; McCarthy, M. K.; Morrison, T. E.; Huey, L.; Annen, K.; DomBourian, M. G.; Knight, V., A New SARS-CoV-2 Dual-Purpose Serology Test: Highly Accurate Infection Tracing and Neutralizing Antibody Response Detection. J Clin Microbiol 2021, 59 (4).

21. Meyer, B.; Reimerink, J.; Torriani, G.; Brouwer, F.; Godeke, G. J.; Yerly, S.; Hoogerwerf, M.; Vuilleumier, N.; Kaiser, L.; Eckerle, I.; Reusken, C., Validation and clinical evaluation of a SARS-CoV-2 surrogate virus neutralisation test (sVNT). Emerg Microbes Infect 2020, 9 (1), 2394–2403.

22. Valcourt, E. J.; Manguiat, K.; Robinson, A.; Chen, J. C.; Dimitrova, K.; Philipson, C.; Lamoureux, L.; McLachlan, E.; Schiffman, Z.; Drebot, M. A.; Wood, H., Evaluation of a commercially-available surrogate virus neutralization test for severe acute respiratory syndrome coronavirus-2 (SARS-CoV-2). Diagn Microbiol Infect Dis 2021, 99 (4), 115294.

23. Alsoussi, W. B.; Turner, J. S.; Case, J. B.; Zhao, H.; Schmitz, A. J.; Zhou, J. Q.; Chen, R. E.; Lei, T.; Rizk, A. A.; McIntire, K. M.; Winkler, E. S.; Fox, J. M.; Kafai, N. M.; Thackray, L. B.; Hassan, A. O.; Amanat, F.; Krammer, F.; Watson, C. T.; Kleinstein, S. H.; Fremont, D. H.; Diamond, M. S.; Ellebedy, A. H., A Potently Neutralizing Antibody Protects Mice against SARS-CoV-2 Infection. J Immunol 2020, 205 (4), 915–922.

24. Chi, X.; Yan, R.; Zhang, J.; Zhang, G.; Zhang, Y.; Hao, M.; Zhang, Z.; Fan, P.; Dong, Y.; Yang, Y.; Chen, Z.; Guo, Y.; Zhang, J.; Li, Y.; Song, X.; Chen, Y.; Xia, L.; Fu, L.; Hou, L.; Xu, J.; Yu, C.; Li, J.; Zhou, Q.; Chen, W., A neutralizing human antibody binds to the N-terminal domain of the Spike protein of SARS-CoV-2. Science 2020, 369 (6504), 650–655.

25. Duan, J.; Yan, X.; Guo, X.; Cao, W.; Han, W.; Qi, C.; Feng, J.; Yang, D.; Gao, G.; Jin, G., A human SARS-CoV neutralizing antibody against epitope on S2 protein. Biochem Biophys Res Commun 2005, 333 (1), 186–93.

26. Muruato, A. E.; Fontes-Garfias, C. R.; Ren, P.; Garcia-Blanco, M. A.; Menachery, V. D.; Xie, X.; Shi, P. Y., A high-throughput neutralizing antibody assay for COVID-19 diagnosis and vaccine evaluation. Nat Commun 2020, 11 (1), 4059.

27. Ye, C.; Chiem, K.; Park, J. G.; Silvas, J. A.; Morales Vasquez, D.; Sourimant, J.; Lin, M. J.; Greninger, A. L.; Plemper, R. K.; Torrelles, J. B.; Kobie, J. J.; Walter, M. R.; de la Torre, J. C.; Martinez-Sobrido, L., Analysis of SARS-CoV-2 infection dynamic in vivo using reporter-expressing viruses. Proc Natl Acad Sci U S A 2021, 118 (41).

28. Schmidt, F.; Weisblum, Y.; Muecksch, F.; Hoffmann, H. H.; Michailidis, E.; Lorenzi, J. C. C.; Mendoza, P.; Rutkowska, M.; Bednarski, E.; Gaebler, C.; Agudelo, M.; Cho, A.; Wang, Z.; Gazumyan, A.; Cipolla, M.; Caskey, M.; Robbiani, D. F.; Nussenzweig, M. C.; Rice, C. M.; Hatziioannou, T.; Bieniasz, P. D., Measuring SARS-CoV-2 neutralizing antibody activity using pseudotyped and chimeric viruses. J Exp Med 2020, 217 (11).

29. Rockx, B.; Kuiken, T.; Herfst, S.; Bestebroer, T.; Lamers, M. M.; Oude Munnink, B. B.; de Meulder, D.; van Amerongen, G.; van den Brand, J.; Okba, N. M. A.; Schipper, D.; van Run, P.; Leijten, L.; Sikkema, R.; Verschoor, E.; Verstrepen, B.; Bogers, W.; Langermans, J.; Drosten, C.; Fentener van Vlissingen, M.; Fouchier, R.; de Swart, R.; Koopmans, M.; Haagmans, B. L., Comparative pathogenesis of COVID-19, MERS, and SARS in a nonhuman primate model. Science 2020, 368 (6494), 1012–1015.

30. Liu, X.; Drelich, A.; Li, W.; Chen, C.; Sun, Z.; Shi, M.; Adams, C.; Mellors, J. W.; Tseng, C. T.; Dimitrov, D. S., Enhanced elicitation of potent neutralizing antibodies by the SARS-CoV-2 spike receptor binding domain Fc fusion protein in mice. Vaccine 2020, 38 (46), 7205–7212.

31. Sha, Y.; Wu, Y.; Cao, Z.; Xu, X.; Wu, W.; Jiang, D.; Mao, X.; Liu, H.; Zhu, Y.; Gong, R.; Li, W., A convenient cell fusion assay for the study of SARS-CoV entry and inhibition. IUBMB Life 2006, 58 (8), 480–6.

32. Yamamoto, M.; Kiso, M.; Sakai-Tagawa, Y.; Iwatsuki-Horimoto, K.; Imai, M.; Takeda, M.; Kinoshita, N.; Ohmagari, N.; Gohda, J.; Semba, K.; Matsuda, Z.; Kawaguchi, Y.; Kawaoka, Y.; Inoue, J. I., The Anticoagulant Nafamostat Potently Inhibits SARS-CoV-2 S Protein-Mediated Fusion in a Cell Fusion Assay System and Viral Infection In Vitro in a Cell-Type-Dependent Manner. Viruses 2020, 12 (6).

33. Wettstein, L.; Weil, T.; Conzelmann, C.; Muller, J. A.; Gross, R.; Hirschenberger, M.; Seidel, A.; Klute, S.; Zech, F.; Prelli Bozzo, C.; Preising, N.; Fois, G.; Lochbaum, R.; Knaff, P. M.; Mailander, V.; Standker, L.; Thal, D. R.; Schumann, C.; Stenger, S.; Kleger, A.; Lochnit, G.; Mayer, B.; Ruiz-Blanco, Y. B.; Hoffmann, M.; Sparrer, K. M. J.; Pohlmann, S.; Sanchez-Garcia, E.; Kirchhoff, F.; Frick, M.; Munch, J., Alpha-1 antitrypsin inhibits TMPRSS2 protease activity and SARS-CoV-2 infection. Nat Commun 2021, 12 (1), 1726.

34. Rajah, M. M.; Bernier, A.; Buchrieser, J.; Schwartz, O., The Mechanism and Consequences of SARS-CoV-2 Spike-Mediated Fusion and Syncytia Formation. J Mol Biol 2022, 434 (6), 167280.

35. Planas, D.; Veyer, D.; Baidaliuk, A.; Staropoli, I.; Guivel-Benhassine, F.; Rajah, M. M.; Planchais, C.; Porrot, F.; Robillard, N.; Puech, J.; Prot, M.; Gallais, F.; Gantner, P.; Velay, A.; Le Guen, J.; Kassis-Chikhani, N.; Edriss, D.; Belec, L.; Seve, A.; Courtellemont, L.; Pere, H.; Hocqueloux, L.; Fafi-Kremer, S.; Prazuck, T.; Mouquet, H.; Bruel, T.; Simon-Loriere, E.; Rey, F. A.; Schwartz, O., Reduced sensitivity of SARS-CoV-2 variant Delta to antibody neutralization. Nature 2021, 596 (7871), 276–280.

36. Rajah, M. M.; Hubert, M.; Bishop, E.; Saunders, N.; Robinot, R.; Grzelak, L.; Planas, D.; Dufloo, J.; Gellenoncourt, S.; Bongers, A.; Zivaljic, M.; Planchais, C.; Guivel-Benhassine, F.; Porrot, F.; Mouquet, H.; Chakrabarti, L. A.; Buchrieser, J.; Schwartz, O., SARS-CoV-2 Alpha, Beta, and Delta variants display enhanced Spike-mediated syncytia formation. EMBO J 2021, 40 (24), e108944.

37. Yurkovetskiy, L.; Wang, X.; Pascal, K. E.; Tomkins-Tinch, C.; Nyalile, T. P.; Wang, Y.; Baum, A.; Diehl, W. E.; Dauphin, A.; Carbone, C.; Veinotte, K.; Egri, S. B.; Schaffner, S. F.; Lemieux, J. E.; Munro, J. B.; Rafique, A.; Barve, A.; Sabeti, P. C.; Kyratsous, C. A.; Dudkina, N. V.; Shen, K.; Luban, J., Structural and Functional Analysis of the D614G SARS-CoV-2 Spike Protein Variant. Cell 2020, 183 (3), 739–751 e8.

38. Zhou, B.; Thao, T. T. N.; Hoffmann, D.; Taddeo, A.; Ebert, N.; Labroussaa, F.; Pohlmann, A.; King, J.; Steiner, S.; Kelly, J. N.; Portmann, J.; Halwe, N. J.; Ulrich, L.; Trueb, B. S.; Fan, X.; Hoffmann, B.; Wang, L.; Thomann, L.; Lin, X.; Stalder, H.; Pozzi, B.; de Brot, S.; Jiang, N.; Cui, D.; Hossain, J.; Wilson, M. M.; Keller, M. W.; Stark, T. J.; Barnes, J. R.; Dijkman, R.; Jores, J.; Benarafa, C.; Wentworth, D. E.; Thiel, V.; Beer, M., SARS-CoV-2 spike D614G change enhances replication and transmission. Nature 2021, 592 (7852), 122–127.

39. Korber, B.; Fischer, W. M.; Gnanakaran, S.; Yoon, H.; Theiler, J.; Abfalterer, W.; Hengartner, N.; Giorgi, E. E.; Bhattacharya, T.; Foley, B.; Hastie, K. M.; Parker, M. D.; Partridge, D. G.; Evans, C. M.; Freeman, T. M.; de Silva, T. I.; Sheffield, C.-G. G.; McDanal, C.; Perez, L. G.; Tang, H.; Moon-Walker, A.; Whelan, S. P.; LaBranche, C. C.; Saphire, E. O.; Montefiori, D. C., Tracking Changes in SARS-CoV-2 Spike: Evidence that D614G Increases Infectivity of the COVID-19 Virus. Cell 2020, 182 (4), 812–827 e19.

40. Ozono, S.; Zhang, Y.; Ode, H.; Sano, K.; Tan, T. S.; Imai, K.; Miyoshi, K.; Kishigami, S.; Ueno, T.; Iwatani, Y.; Suzuki, T.; Tokunaga, K., SARS-CoV-2 D614G spike mutation increases entry efficiency with enhanced ACE2-binding affinity. Nat Commun 2021, 12 (1), 848.

41. Jiang, X.; Zhang, Z.; Wang, C.; Ren, H.; Gao, L.; Peng, H.; Niu, Z.; Ren, H.; Huang, H.; Sun, Q., Bimodular effects of D614G mutation on the spike glycoprotein of SARS-CoV-2 enhance protein processing, membrane fusion, and viral infectivity. Signal Transduct Target Ther 2020, 5 (1), 268.

42. Mlcochova, P.; Kemp, S. A.; Dhar, M. S.; Papa, G.; Meng, B.; Ferreira, I.; Datir, R.; Collier, D. A.; Albecka, A.; Singh, S.; Pandey, R.; Brown, J.; Zhou, J.; Goonawardane, N.; Mishra, S.; Whittaker, C.; Mellan, T.; Marwal, R.; Datta, M.; Sengupta, S.; Ponnusamy, K.; Radhakrishnan, V. S.; Abdullahi, A.; Charles, O.; Chattopadhyay, P.; Devi, P.; Caputo, D.; Peacock, T.; Wattal, C.; Goel, N.; Satwik, A.; Vaishya, R.; Agarwal, M.; Indian, S.-C.-G. C.; Genotype to Phenotype Japan, C.; Collaboration, C.-N. B. C.-.; Mavousian, A.; Lee, J. H.; Bassi, J.; Silacci-Fegni, C.; Saliba, C.; Pinto, D.; Irie, T.; Yoshida, I.; Hamilton, W. L.; Sato, K.; Bhatt, S.; Flaxman, S.; James, L. C.; Corti, D.; Piccoli, L.; Barclay, W. S.; Rakshit, P.; Agrawal, A.; Gupta, R. K., SARS-CoV-2 B.1.617.2 Delta variant replication and immune evasion. Nature 2021, 599 (7883), 114–119.

43. Saito, A.; Irie, T.; Suzuki, R.; Maemura, T.; Nasser, H.; Uriu, K.; Kosugi, Y.; Shirakawa, K.; Sadamasu, K.; Kimura, I.; Ito, J.; Wu, J.; Iwatsuki-Horimoto, K.; Ito, M.; Yamayoshi, S.; Loeber, S.; Tsuda, M.; Wang, L.; Ozono, S.; Butlertanaka, E. P.; Tanaka, Y. L.; Shimizu, R.; Shimizu, K.; Yoshimatsu, K.; Kawabata, R.; Sakaguchi, T.; Tokunaga, K.; Yoshida, I.; Asakura, H.; Nagashima, M.; Kazuma, Y.; Nomura, R.; Horisawa, Y.; Yoshimura, K.; Takaori-Kondo, A.; Imai, M.; Genotype to Phenotype Japan, C.; Tanaka, S.; Nakagawa, S.; Ikeda, T.; Fukuhara, T.; Kawaoka, Y.; Sato, K., Enhanced fusogenicity and pathogenicity of SARS-CoV-2 Delta P681R mutation. Nature 2022, 602 (7896), 300–306.

